# GABA relates to functional connectivity changes and retention in visuomotor adaptation

**DOI:** 10.1101/2020.12.22.423981

**Authors:** Caroline Nettekoven, Sinead Brady, William T Clarke, Uzay Emir, Jacob Levenstein, Pierre Petitet, Muriel T N Panouilleres, Velicia Bachtiar, Jacinta O’Shea, Heidi Johansen-Berg, Ned Jenkinson, Charlotte Stagg

## Abstract

Motor adaptation is fundamental to maintaining accurate movements under changing conditions, but the neurochemical basis of human motor adaptation is not well understood. Here, we used an ultra-high-field MR multimodal acquisition to address the hypothesis that M1 GABA and M1-Cerebellar functional connectivity would relate to retention of adaptation, but not acquisition of adaptation. As such, we demonstrate higher baseline M1 [GABA] relates to greater retention but does not relate to adaptation-acquisition. This relationship is mediated by change in M1-Cerebellar functional connectivity: higher M1 [GABA] relates to a decreased M1-Cerebellar connectivity, resulting in greater retention. These findings showed anatomical, neurochemical and behavioural specificity: As expected, no relationship was found between retention and a control metabolite, retention and connectivity change between control regions and between M1 [GABA] and behaviour in a control condition. The implication of a mechanistic link from neurochemistry to retention advances our understanding of population variability in retention and provides a step towards therapeutic interventions to restore motor abilities.

## Introduction

Interacting with our ever-changing physical environment requires continual recalibration of the motor system. One mechanism by which this occurs is motor adaptation. Understanding how motor adaptation is implemented by the human brain, how different regions work in concert to retain adaptive movement accuracy, and how this function is linked to metabolic use of neurochemicals poses an important challenge in neuroscience. Transcranial direct current stimulation (tDCS) data has suggested that acquisition of motor adaptation is reliant on the cerebellum but not M1, while storage of the adapted state (retention) may depend on M1 {Galea2011} (but see {Oldrati2018}{Jalali2018}). Magnetic Resonance Spectroscopy (MRS) data has shown that M1 GABA concentration predicts behaviour that depends on M1 {Kolasinski2019}. Therefore, we hypothesised that in adaptation, where behaviour is acquired outside M1, but retained within M1, GABAergic tone in M1 would correlate with retention but show no relationship with acquisition of adaptation. Further, adaptation has been shown to change connectivity between the cerebellum and M1 {Mawase2017}. We therefore hypothesised that the link between M1 GABA and retention would be mediated by a change in connectivity between M1 and cerebellum. To test these hypotheses, we quantified [GABA] and [Glutamate] from the hand region of the left human primary motor cortex (M1) using ultra-high-field magnetic resonance spectroscopy (7T-MRS) while participants (n=15; mean age 26 ± 3 years, range 22-32 years) performed a visuomotor task with or without an adaptation component (cursor rotation condition vs. control condition) in a within-subject, crossover study. To assess connectivity change, we collected resting-state fMRI data immediately before and after the task. To probe retention participants performed a washout behavioural task after each MR session (Fig. 1A).

**Figure 1.**
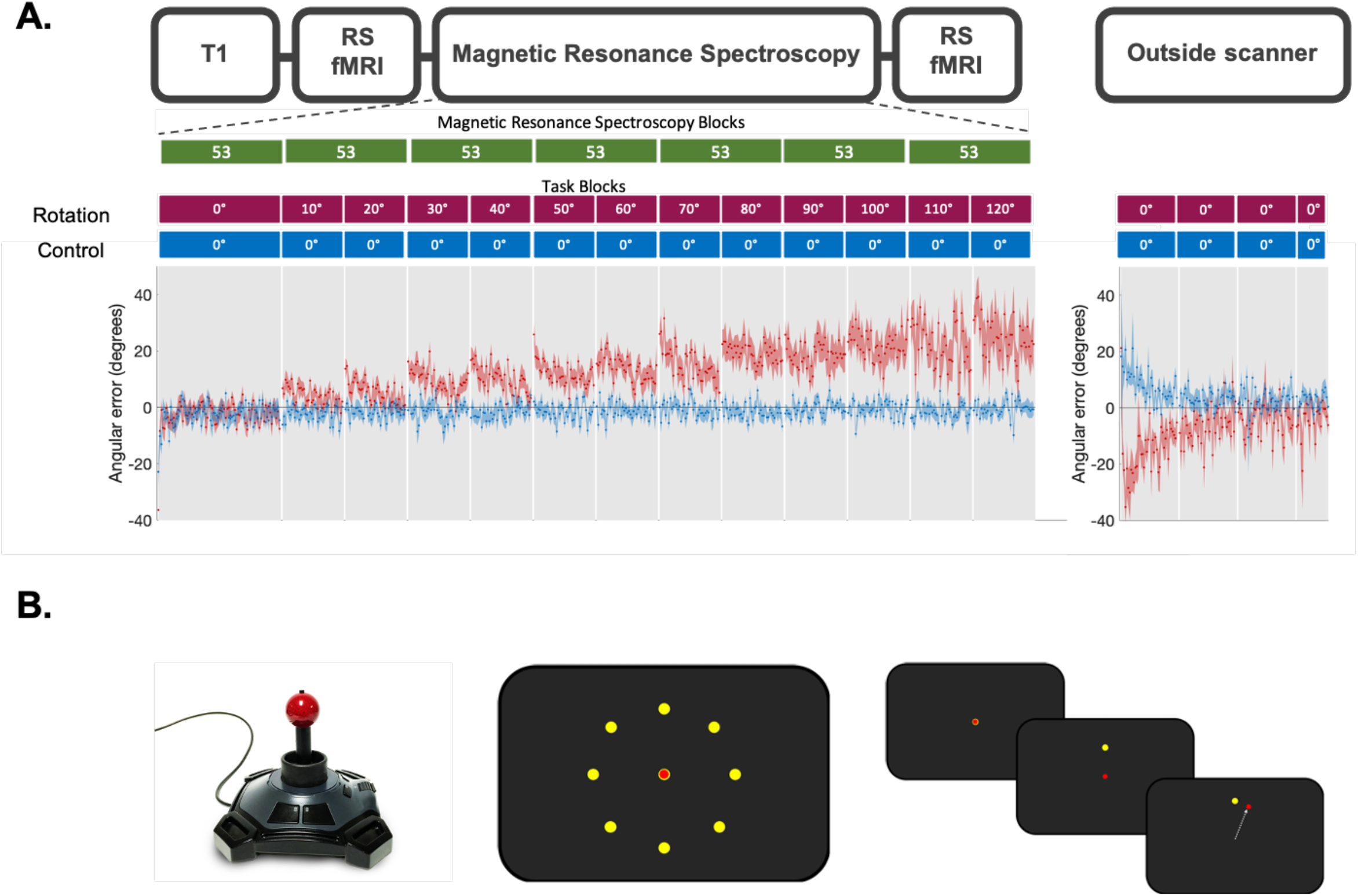
Experiment. **A.** Experiment Protocol and Behaviour. **Upper panel.** Scanning Protocol. T1-weighted structural image was acquired at the begin of the MRI scan. Resting-state fMRI data was acquired before and after the task. Magnetic Resonance Spectroscopy (MRS) data was acquired during the task. Each MRS-block consisted of 53 individual spectra (MRS-blocks shown in green), which were averaged. This resulted in seven MRS-blocks. **Middle Panel.** Task Protocol. In the rotation condition (red blocks), stepwise increasing rotated visual feedback required participants to adapt movements to reduce errors. One block at each angle and each block consisted of 40 trials of 4 seconds duration each. The numbers in the red and blue boxes indicate the degree to which the visual feedback was rotated, with 0° indicating no rotation. In the control session (blue blocks), participants performed the task without any rotation imposed, but for the same length (560 trials in total for the main task). The rotation was washed out after task (144 trials, no rotation). The task was practised before the main task outside of the scanner and inside of the scanner (32 trials each, no rotation; not depicted in this figure but see supplementary figure 1.). **Lower panel.** Behavioural data. Shown is angular error at each trial averaged across participants. Shaded area represents standard error of the mean. Rotation condition error is shown in red. Control condition error is shown in blue. **B.** Task Schematic. **Left.** MR-compatible Joystick used to record participant responses. **Middle.** Eight possible target locations (yellow) centred radially around the cursor (red) at its starting position. **Right.** Schematic of a rotation trial. Cursor (red) is first presented at the centre starting position. Target (yellow) appears at one of the eight possible target locations. Participant makes a centre-out movement towards the target but sees clockwise rotated visual feedback.

## Results

### Participants were able both to adapt to the perturbation and to retain the adapted state

In the rotation condition, participants were required to adapt their centrifugal shooting movements to a rotation of the visual feedback which increased stepwise by 10 degrees after every block of 40 trials in order to maximally drive adaptation throughout the duration of the scanning session. Trials were grouped into epochs, with one epoch containing eight consecutive trials and all blocks containing five epochs. All participants were able to learn the task: we observed a significant reduction of error within the adaptation blocks from the first to the last epoch, indicating that participants adapted (Fig.1A, left panel). This finding was confirmed by a significant effect of epoch (χ2(1) = 55.597, p < 0.0001) in a linear mixed effect model of error in the first and last epoch across all blocks. There was no such change in error in the control condition (χ2(1) = 0.7, p = 0.4), suggesting that, as hypothesised, participants adapted their movements only in the rotation condition. This was confirmed by a direct comparison between error reduction in the rotation condition and error reduction in the control condition, showing that participants adapted significantly more in the rotation condition (paired t-test between error reduction in rotation condition and in control condition, with error reduction calculated as the mean difference between error in first epoch and last epoch of all blocks after baseline: t(11) = 3.69, p < 0.05). Post-hoc t-tests on each separate block of the rotation condition showed that participants were able to adapt to rotations up to 70° (paired t-test of first and last epoch: t(12) > 3.58, p < 0.05, Bonferroni adjusted), but not above 70° (Five t-tests, one for each block with a rotation greater than 70°, all t(12)< 0.62, p > 0.5). We therefore excluded blocks with a rotation of 80° or more when calculating our adaption measure. The adaptation measure was therefore quantified as the mean angular error during the rotation condition across rotation blocks up to 70°. We also excluded the final two MRS-blocks from further analysis for the same reason.

Participants retained the previously learned adaptive movement in the first block of the washout (one-sample t-test of error in first block of washout in rotation condition: t(12) = −9.635, p < 0.00001; red line in Fig.1A right panel), with retention assessed approximately 22 minutes after adaptation acquisition (mean time gap of 22.43 ± 6.56 minutes). Participants also retained a fraction of the compensatory movement in the second and third block of the washout (Second: t(12) > −4.108, p < 0.01; Third: t(12) > −9.63, p < 0.05, Bonferroni adjusted). A per-subject measure of retention was quantified from the angular error in the first block of the washout after adaptation, since participants showed the most retention in this first block (mean angular error of −13.68 ± 5.12). We calculated retention as mean angular error across all trials in the first washout block. We found no correlation between adaptation and retention on a subject-by-subject basis (r = −0.01, p = 0.75), even when controlling for any differences in the time elapsed (r = −0.13, p = 0.67). In the control condition, participants also showed a positive mean angular error during the first block of the washout (one-sample t-test: t(12) = 7.82, p < 0.01; blue line in Fig.1A right panel). Visual inspection of the 32 practice trials performed inside the scanner before the main task revealed that this positive error in the control condition was likely driven by an unspecific movement bias due to the participants position inside the scanner (Supplementary Fig. 1).

### Adaptation led to an increase in connectivity in the cerebellar network

We designed the adaptation task used in this study to maximise the amount of time that cerebellar-dependent mechanisms of adaptation could be driven, by imposing a rotation of the visual feedback that increased by 10 degrees every 40 trials. However, this meant the adaptation paradigm was unusual in its use of a stepwise increasing rotation. To ensure this particular adaptation paradigm drove plastic changes known to occur in response to adaptation, we specifically set out to replicate the finding that adaptation modulates connectivity of a cerebellar resting-state network, compared to simple movement execution {Albert2009}. As expected, the cerebellar network significantly increased in strength after performing the adaptation task (cerebellar regions that increased in network connectivity shown in red-yellow in Fig. 2A). Moreover, we hypothesised that in the rotation condition, change in strength of functional connectivity within the cerebellar network correlates with adaptation, such that participants who show an increase in cerebellar network strength would adapt better, which goes beyond the findings of Albert and colleagues. To test this, we calculated change in strength of cerebellar network for each subject in the rotation condition by subtracting pre network strength from post network strength. We also quantified adaptation error for each subject by calculating the mean error across all rotation blocks in which adaptation took place, excluding the first epoch of each block. Indeed, there was a correlation between cerebellar network change and adaptation error (r = −0.648, p = 0.016; Fig. 2C), such that participants who adapted better also showed an increase in strength of the cerebellar network. This relationship was behaviourally specific to adaptation (correlation between network strength change and retention, r = −0.038, p = 0.9, zdiff (adaptation vs retention) = −1.71, p = 0.0439; correlation between network strength change and control behaviour r = 0.138 p = 0.64, zdiff (adaptation vs control) = −2.17, p = 0.015). To test network specificity, we ran the same analyses on the default mode network (DMN), a well-described network, and saw no change in network strength (no increases in network connectivity, Fig. 2B), nor a significant correlation of change in DMN strength with adaptation (Fig. 2D; r = 0.34, p = 0.28, zdiff (DMN vs cerebellar network) = −2.61, p = 0.0045).

**Figure 2.**
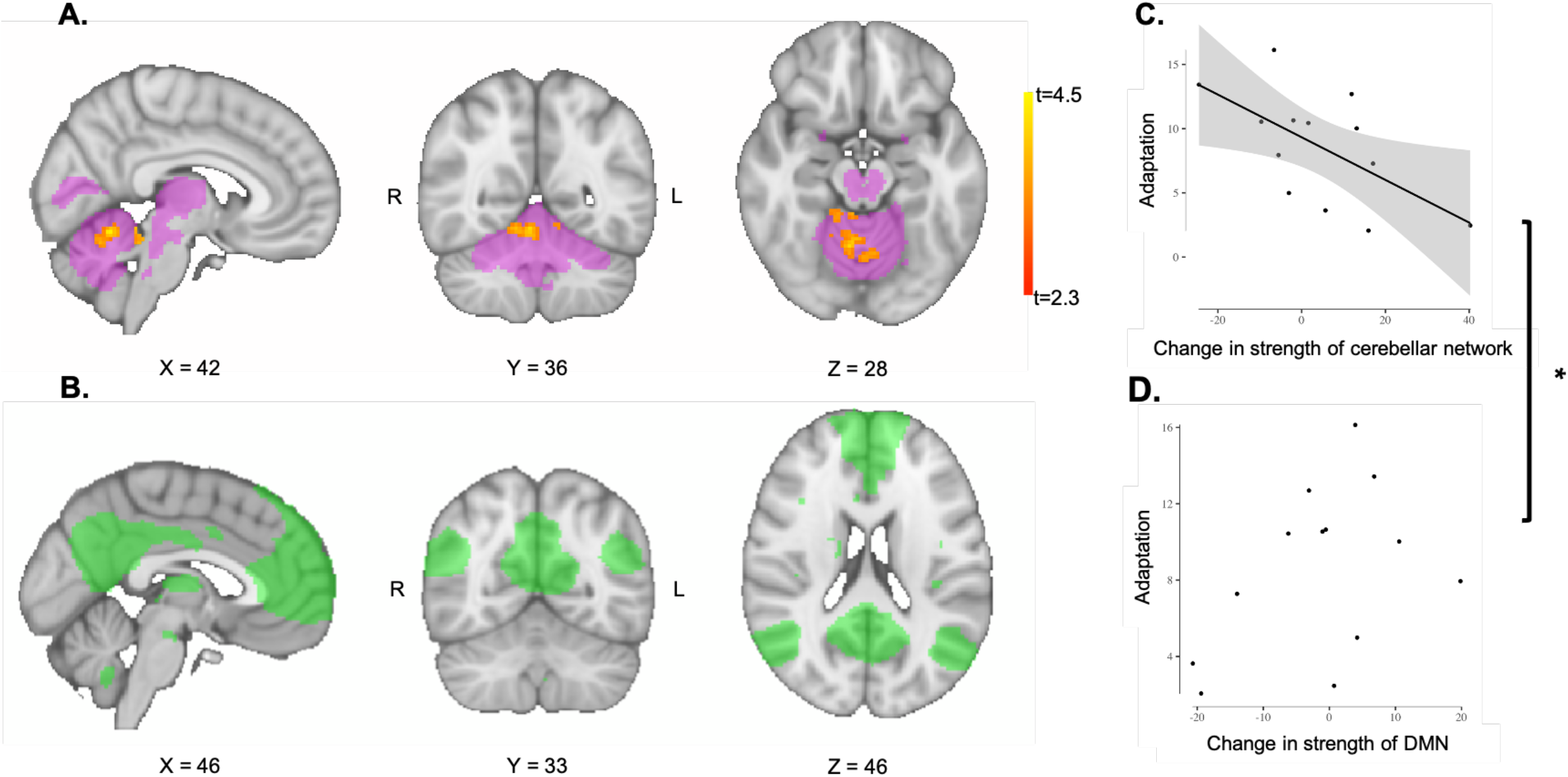
Adaptation increased functional connectivity in cerebellar network in a highly specific and performance-relevant manner. **A.** Functional connectivity increases in cerebellar network after adaptation. Cerebellar network is shown in pink. Voxels that show increased functional connectivity after performance of the rotation task compared toafter performance of the control task are shown in red-yellow (Rotation vs. Control in post network). Colour bar indicates range of t-values in those voxels that are significant (family-wise error controlled, thresholded at p < 0.05). Brain slices are shown according to radiology convention, so left hemisphere is shown on the right (indicated by R and L surrounding the coronal slice). **B.** No change in functional connectivity of Default Mode Network. The Default Mode Network (DMN) was chosen as a control network and is shown in green. As expected, adaptation did not affect functional connectivity in the DMN (all p > 0.24). **C.** Change in cerebellar network strength correlates with adaptation performance. Participants who adapted better to the rotation, show an increase in network strength from before the rotation task performance to after (r = −0.648, p = 0.016). Participants who adapted less, as shown by a larger adaptation error, show a decrease in network strength from pre to post, as shown by negative values. Datapoints are individual participants. Plot shows line of best fit for correlation and 95% confidence interval. **D.** Change in Default Mode Network strength does not correlate with adaptation performance. No relationship between change in DMN network strength from before to after rotation task performance and adaptation error (r = 0.34, p = 0.28). Datapoints are individual participants. Line of best fit is not plotted for this correlation, because the correlation is not significant. * Indicates significant difference between correlations shown in C and in D: The correlation coefficient of Cerebellar network change and adaptation error was significantly different from the correlation coefficient of DMN change and adaptation error (zdiff = −2.61, p = 0.0045)

### [GABA] in M1 relates to adaptation retention, but not adaptation acquisition

Next, we wished to investigate the role of M1 [GABA] in adaptation and retention. In this within-subject study, the M1 MRS voxel was reproducibly placed between sessions (mean spatial correlation coefficient between each subject’s pair of MRS voxels: r = 0.62 ± 0.38; Fig. 3A) and coverage of the left M1 hand region did not differ between sessions (t(11) = 0.37, p = 0.72; Fig. 3A).

**Figure 3.**
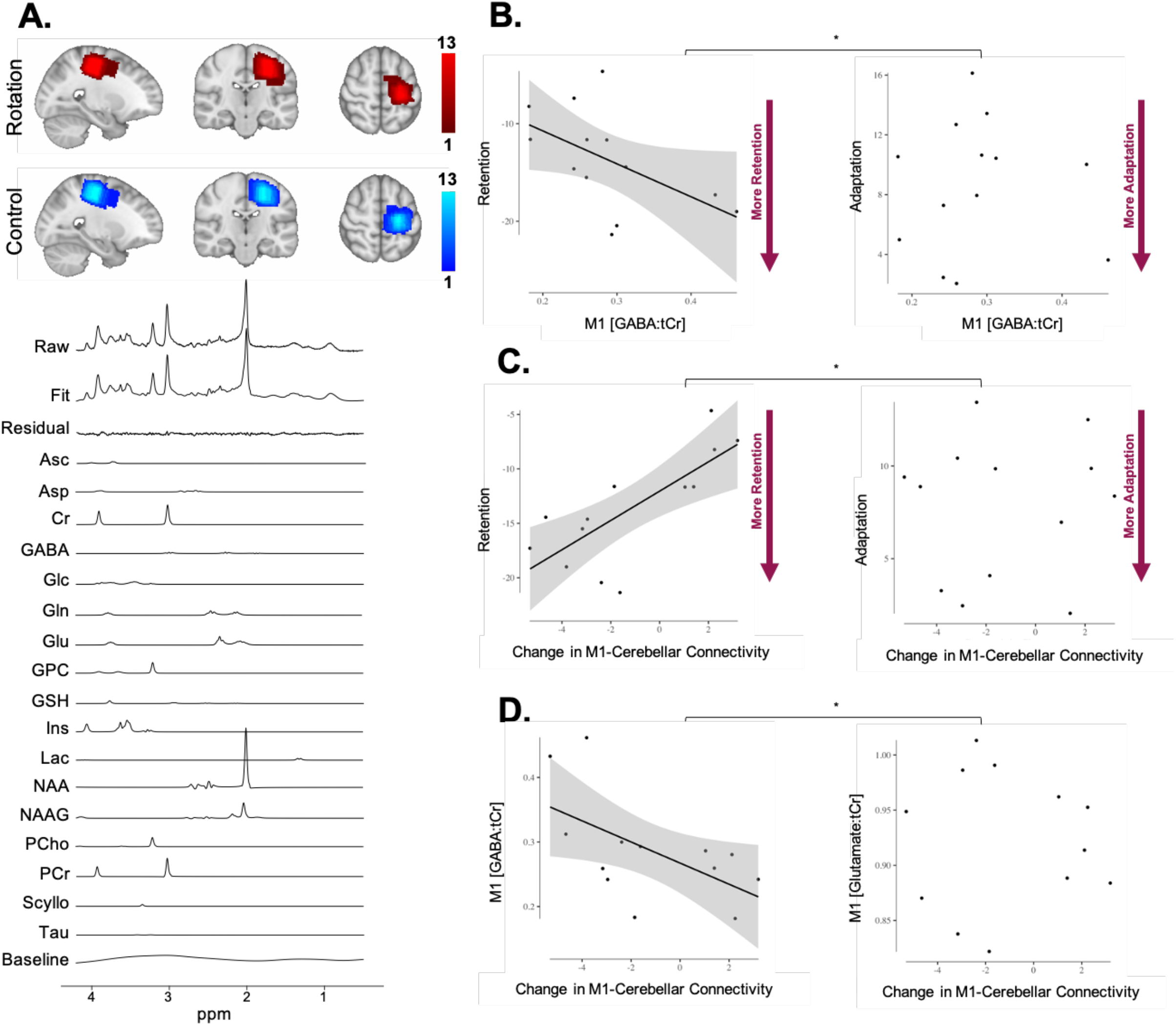
M1 GABA correlates with retention and M1-Cerebellar connectivity change. **A.** Voxel Placement and Spectral Fit. **Upper panel.** MRS voxel overlap maps for the two conditions. 20 x 20 x 20 mm3 voxel centred over hand area of left M1. Colour bars represent number of participants. **Lower panel.** Spectral Fit. Image shows representative spectra of one subject including LCModel fit. Metabolite basis spectra have been scaled using LCModel such that a linear combination of the basis spectra, the residual and the baseline best fits the raw measured spectrum. Measured raw spectrum is shown in the at the top and LCModel Fit is shown below. GABA metabolite peaks appear at 1.89 ppm, 2.29 ppm and 3.01 ppm. **B, C & D.** Each dot is one participant. Due to technical failures during MRI, two participants were excluded from analysis. Shading indicates 95% confidence bounds of the relationship. **B.** M1 [GABA] correlates with retention, but not adaptation. **Left.** Baseline M1 GABA was significantly correlated with retention, with higher concentration of M1 GABA associated with greater retention (r(13) = −0.62 p = 0.02). **Right.** Baseline M1 GABA and adaptation did not correlate (r(13) = 0.07 p = 0.82). The correlations between M1 GABA and retention (left panel) and M1 GABA and adaptation (right panel) were significantly different (zdiff = −1.87, p = 0.03). **C.** Change in M1-Cerebellar connectivity correlates with retention, but not adaptation. **Left.** M1-Cerebellar connectivity change and retention were significantly correlated (r(13) = 0.68, p = 0.01). A decrease in M1-Cerebellar connectivity was linked to greater retention. **Right.** M1-Cerebellar connectivity change and adaptation did not correlate (r(13) = 0.05, p = 0.87. The correlation between M1-Cerebellar connectivity change and retention (left panel) and M1-Cerebellar connectivity change and adaptation (right panel) were significantly different (zdiff (retention vs adaptation) = 1.73, p = 0.04). **D.** Change in M1-Cerebellar connectivity correlates with correlates with M1 [GABA], but not M1 [Glutamate]. **Left.** Baseline M1 GABA and M1-Cerebellar connectivity change were significantly correlated. Higher baseline [GABA] in left M1 was associated with a greater subsequent decrease in connectivity between left M1 with the right cerebellum (r(13) = −0.63, p = 0.03). **Right.** Baseline M1 Glutamate and M1-Cerebellar connectivity change did not correlate (r(13) = −0.08, p = 0.8). The correlation between M1 GABA and M1-Cerebellar connectivity change (left panel) and M1 Glutamate and M1-Cerebellar connectivity change (right panel) were significantly different (zdiff (GABA vs Glutamate) = −1.81, p = 0.03).

To test whether GABA changes during adaptation, we constructed a linear mixed effects model of GABA with a fixed effect of MRS-block (1-5) and a fixed effect of condition (rotation, control). Consistent with our hypothesis that adaptation relies on brain regions distant from M1, we saw no change in GABA in the rotation condition compared to the control condition (no significant MRS-block x condition interaction: χ2(1) = 3.32, p = 0.34; main effect of MRS-block χ2(3) = 1.66, p = 0.65; main effect of condition χ2(1) = 0.31, p = 0.58). There was no relationship between baseline M1 [GABA] and our behavioural measure of adaptation (r(13) = 0.07 p = 0.82; Fig. 3B right panel). However, in line with the hypothesis that retention is M1-dependent, baseline M1 [GABA] was significantly correlated with retention, with higher concentration of M1 [GABA] associated with greater retention (r(13) = −0.62 p = 0.02; Fig. 3B left panel). The correlations between M1 [GABA] and retention and M1 [GABA] and adaptation were significantly different (zdiff = −1.87, p = 0.03). To ensure the robustness of the observed correlation between M1 [GABA] and retention against potential outliers, we used robust correlation methods selected for its suitability to the given data structure according to criteria outlined in {Pernet2013} to re-calculate the correlation coefficient. The robust correlation method confirmed a significant relationship between M1 [GABA] and retention (percentage-bend correlation coefficient r(13) = 0.63 CI = [−0.87 −0.19]).

To determine the specificity of the link between M1 [GABA] and retention, we tested whether the same relationship existed for a control neurotransmitter (glutamate) and within the control behavioural condition. The correlation between retention and baseline M1 [Glutamate] did not reach significance (r(13) = −0.57 p = 0.0546, no significant difference to correlation coefficient of M1 [GABA] and retention: zdiff = −0.19, p = 0.42). There was no relationship between the retention error equivalent of the control condition and baseline M1 [GABA] in the control condition (r(12) = −0.02 p = 0.9574, difference to correlation coefficient of M1 [GABA] and retention: zdiff = −1.74, p = 0.04).

### Connectivity between M1 and cerebellum relates to retention

We next reasoned that if adaptation occurs in the cerebellum and retention is M1 dependent, then the change in connectivity between these two regions would likely be related to the degree of retention. M1-Cerebellar connectivity was calculated by cross-correlating the mean BOLD time course from the hand region in left M1 with the time course from a right cerebellar region involved in adaptation{Hardwick2013}. Change in connectivity was calculated by subtracting the correlation coefficient of the extracted timeseries in the pre-scan from the correlation coefficient in the post-scan (i.e. post minus pre, yielding positive values when connectivity increases). We observed a significant relationship between M1-Cerebellar connectivity change and retention (r(13) = 0.68, p = 0.01; Fig. 3C, left panel), indicating that a decrease in M1-Cerebellar connectivity was linked to greater retention. Robust correlation methods confirmed this significant relationship (percentage-bend correlation: r(13) = 0.78 CI=[0.12 0.94], {Wilcox1994}{Pernet2013}). The relationship between connectivity change and retention was also behaviourally-specific (no significant correlation between M1-Cerebellar connectivity change and adaptation (r(13) = 0.05, p = 0.87; zdiff (retention vs adaptation) = 1.73, p = 0.04 Fig. 3C right panel).

Supporting the hypothesis that M1 [GABA] is associated with change in connectivity, there was a significant correlation between baseline M1 [GABA] and M1-Cerebellar connectivity change, such that higher baseline [GABA] in left M1 was associated with a greater subsequent decrease in connectivity between left M1 with the right cerebellum (r(13) = −0.63, p = 0.03; Fig. 3D left panel). This significant relationship was confirmed by robust correlation methods (percentage-bend correlation coefficient r(13) = −0.55 CI = [−0.87 −0.02]).

To determine the specificity of the link between M1 [GABA], connectivity and retention, we tested whether the same relationships existed for a control neurotransmitter (glutamate), a control anatomical region (left cerebellum) and a control behavioural condition (control “retention”). There was no correlation between glutamate and M1-right Cerebellar connectivity change (r(13) = −0.08, p = 0.8; zdiff (GABA vs Glutamate) = −1.81, p = 0.03; Fig. 3D right panel), nor between M1 [GABA] and connectivity change between M1 and left cerebellum (r(13) = −0.37, p = 0.21, zdiff = −0.64, p = 0.26). There was no correlation between retention and connectivity change between left M1 and left cerebellum (r(13) = 0.38 p = 0.19; ⍰zdiff = 0.93, p = 0.18). We observed no correlation between control “retention” and M1-Cerebellar connectivity in the control condition (r(14) = 0.04, p = 0.89; zdiff = 1.78 p = 0.04).

### Connectivity change mediates the relationship between M1 [GABA] and retention

In order to explore the potential role of change in connectivity as a mechanism through which baseline M1 [GABA] could influence later retention, we performed a mediation analysis (Fig. 4). A mediation analysis tests whether the observed relationship between M1 [GABA] and Retention (path c) might be statistically mediated through an indirect pathway (via M1–cerebellar connectivity change; path ab). As expected, M1 [GABA] significantly related to both retention (path c = −33.74, p = 0.0005) and M1-Cerebellar Connectivity Change (path a = −20.14, p = 0.02). M1-Cerebellar Connectivity Change in turn was associated with retention (path b = 1.19, p = 0.017). The relationship between M1 [GABA] and retention was mediated by connectivity change with a 71% percentage mediation (ab = −23.87; CI[−62.29, −1.80]). Once M1-Cerebellar Connectivity Change was added to the model, the direct effect of M1 [GABA] on retention became non-significant (c’ = −9.87, p = 0.39), meaning the relationship is fully mediated. Together, these findings suggest that M1-Cerebellar connectivity change might be a potential mechanism through which baseline M1 [GABA] influences later retention.

**Figure 4.**
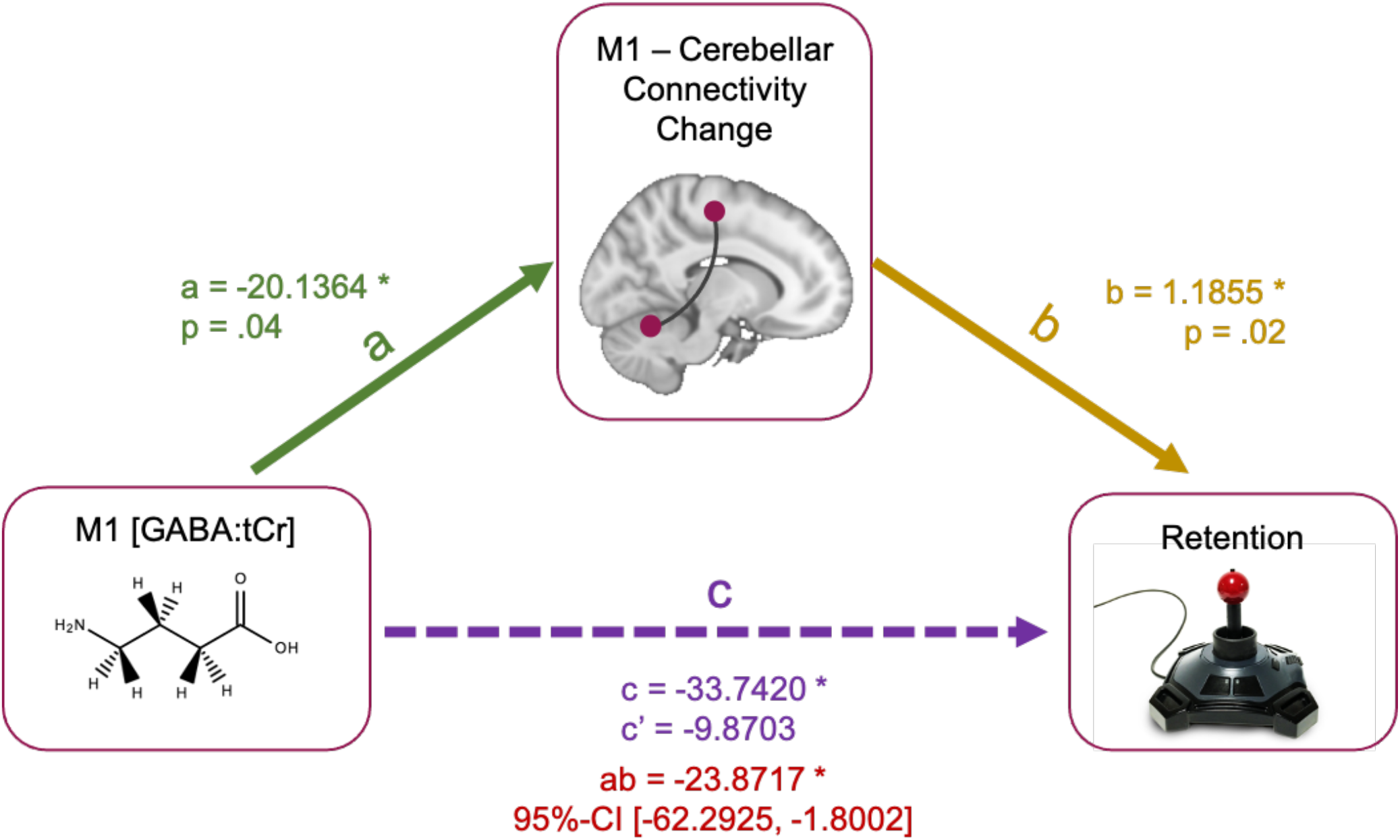
M1-Cerebellar connectivity change mediates the relationship between M1 [GABA] and retention. A mediation analysis tests whether the observed direct relationship between two variables, here M1 [GABA] and Retention (path c; dotted arrow), might be mediated through an indirect pathway, here M1–cerebellar connectivity change (path ab; continuous arrows). The numbers are the coefficients of each path. * Indicates significant coefficients. M1 GABA significantly predicted both retention (path c = −33.74, p = 0.0005) and M1-Cerebellar connectivity change (path a = −20.14, p = 0.02). M1-Cerebellar connectivity change in turn predicted retention (path b = 1.19, p = 0.017). The relationship between M1 GABA and retention was mediated by connectivity change with a 71% percentage mediation (ab = −23.87; 95%-CI[−62.2925, −1.8002]). Once M1-Cerebellar connectivity change was added to the model, the direct effect of M1 [GABA] on retention became non-significant (c’= −9.87, p = 0.39), meaning the relationship is fully mediated.

## Discussion

This study provides evidence that retention of a visuomotor transformation may be explained by the underlying neurochemical processes and functional changes in the human brain. Specifically, higher M1 [GABA] concentration before acquisition of a visuomotor adaptation is associated with a greater decrease in M1-Cerebellar connectivity during acquisition, and this, in turn, is associated with more retention of the adapted state afterwards. These relationships are anatomically specific, since retention of the right-handed adaptation task can be related only to connectivity change between contralateral M1 and ipsilateral cerebellum and not to connectivity change between contralateral M1 and contralateral cerebellum. The demonstrated relationships are also behaviourally specific, since neurochemical and functional differences relate to retention, but not acquisition of the transformation. Finally, the relationship is neurochemically specific, as glutamate concentration does not significantly correlate with retention. To ensure our findings are robust against potential outliers, we re-calculated all significant relationships using robust correlation methods, resulting in the same pattern of results. We observed a positive angular error in the washout phase of the control condition. This control washout error seems to be driven by the positioning of the participant inside the scanner: Inspection of the practice trials performed inside and outside of the scanner (practice 1 and practice 2 in Supplementary Fig. 1A, respectively) revealed a movement bias as soon as the participant is placed into the scanner. This bias reduces to zero within few trials and the inverse of the bias matches the control condition washout error (Supplementary Fig. 1B), suggesting that the scanner position drives the control washout error. We observed no significant correlation between neurochemical and functional differences and this movement bias in the control condition, suggesting that the demonstrated relationships are specific to the rotation condition.

Motor adaptation is a form of motor learning where a transformation of motor commands is required to maintain accurate movements after a change in the environment or in the body. Crucial sites for adaptation include the cerebellum and M1 and these regions have been purported to underly different aspects of motor adaptation. There is emerging evidence that M1 may be involved in adaptation of distal limb movements (for example finger movements) {Weightman2020} {Panouilleres2015}. However, for tasks involving more proximal effectors (for example gross upper arm movements), acquisition in motor adaptation predominantly involves the cerebellum {Panouilleres2015a} {Miall2005}, whereas retention of the adapted state is classically believed to rely on M1 {Herzfeld2014} {Hunter2009} {Richardson2006}. To this effect, increasing M1 excitability with anodal tDCS facilitates retention, but does not influence adaptation {Galea2011}, and disrupting M1 activity with TMS impairs retention without affecting adaptation {Hadipour-Niktarash2007}. Previous work has shown an association between M1 [GABA] change induced by tDCS during prism adaptation and change in retention induced by tDCS {OShea2017} {Petitet2018}, with a larger decrease in M1 [GABA] relating to greater retention change. Further, retention of prism adaptation has been related to baseline M1 [GABA] such that participants with lower M1 [GABA] retained more of the adapted state {Petitet2018}, overall suggesting that lower M1 [GABA] facilitates retention. Conversely, outside of the adaptation domain, retention of memories has been linked to an increase in visual area [GABA:Glu] ratio for visual perception learning {Shibata2017; Tamaki2020} and an increase in hippocampal [GABA:Glu] ratio for spatial learning {Koolschijn2019}, suggesting that more GABA-dominant processing protects against memory interference and facilitates retention. We find that in visuomotor adaptation, higher M1 [GABA] at baseline is associated with greater subsequent retention, in line with a facilitative effect of GABA for retention. The contrasting results in retaining prism adaptation and visuomotor rotation adaptation may be explained by the two paradigms eliciting different contributions of adaptation processes: simultaneous, but distinct processes have been shown to underlie adaptation {Smith2006} {Taylor2014} {Taylor2011} {McDougle2015}, and the adaptation paradigm has been suggested to influence which processes are favoured {Fleury2019}, which may in turn result in differential recruitment of M1. Alternatively, differential involvement of M1 during the task may also be driven by the requirement to adapt proximal limb movements for prism adaptation and distal limb movements for the visuomotor adaptation task presented here. However, the extent to which the choice of adaptation paradigm recruits different circuits in the brain is currently unclear.

Our data are consistent with the proposed role of M1 in retention but not adaptation, and further suggest that connectivity change between M1 and the cerebellum may reflect the means by which the adapted state, once acquired is retained. These results are in line with evidence from locomotor adaptation studies that have investigated changes in M1-Cerebellar interactions after acquisition of the novel locomotor transformation. Specifically, in locomotor adaptation, connectivity between M1 and the cerebellum decreased {Mawase2017}, and the inhibitory tone of the cerebellum over M1 assessed with paired pulse TMS (cerebellar-brain inhibition (CBI); {Ugawa1995}, Pinto and Chen 2001) decreased after acquisition of the locomotor transformation {Jayaram2011}. Moreover, the latter study found that the extent of CBI reduction correlated with retention, such that participants who showed greater reduction retained more {Jayaram2011}.

Here, we demonstrate that the baseline [GABA] in M1 is associated with change in functional connectivity between the cerebellum and M1 and retention of behaviour. While the physiological basis of MRS-assessed GABA has yet to be fully elucidated, it has been proposed to primarily reflect extracellular tonic GABA signalling {Semyanov2004}, rather than phasic, synaptic signalling {Dyke2017} {Stagg2011a}, though physiologically these two forms of inhibition are related {Farrant2005}. MRS-assessed GABA has been linked to oscillatory activity in the gamma range {Towers2004}, which in turn has been shown to relate to long-range connectivity {Cabral2011} {Shmuel2008}. This hypothesis is supported by previous work from our lab and others, suggesting that [GABA] in a key network node is related to functional connectivity within the functional network {Stagg2014} {Bachtiar2015} {Kapogiannis2013}. Our finding that greater decreases in M1-Cerebellar connectivity were observed in participants with higher baseline M1 [GABA] is therefore in keeping with this mechanistic framework. Further, since a decrease in connectivity is commonly interpreted as indicating a segregation of processing between two brain regions, a plausible interpretation of our results would be that higher M1 [GABA] before adaptation leads to more segregated processing of motor information during adaptation. This segregation of processing, in turn, may result in better subsequent retention of the visuomotor transformation, though these hypotheses remain to be tested.

The results presented here demonstrate a series of dependent integrated mechanisms from local neurochemistry, to functional connectivity between brain regions, to retention of adapted behaviours. These findings not only significantly advance our understanding of population variability in retention behaviour, but also, in light of the therapeutic potential of interventions that drive retention to mitigate treatment-resistant effects of stroke {OShea2017}, they also provide a crucial step towards developing therapeutic interventions to restore motor abilities.

## EXPERIMENTAL MODEL AND SUBJECT DETAILS

15 healthy, right-handed participants (8 female; mean age 26 years, range 22-32 years) gave their written informed consent in line with local ethics committee approval (Oxford CUREC C1-2014-090). Each participant had two MR scans while performing a visuomotor task that either involved a learning component (adaptation task) or no learning component (control task). The order of sessions was counterbalanced across the group. Participants were right-handed, had no history of neurological or psychiatric conditions, were not taking any medications that can affect the central nervous system, and had no contraindications to MRI. Due to technical failures during MRI, two participants were excluded from analysis.

## METHOD DETAILS

### Behavioural data acquisition

Participants controlled a cursor on a screen using a joystick and made discrete movements to a target. They were instructed to make fast, accurate and ballistic movements to ‘shoot’ through the target that appeared on the screen. The target appeared at one of eight locations radially aligned around the starting position of the cursor and separated by 45° (Fig. 1B). Trials were grouped into epochs and blocks, with one epoch containing eight consecutive trials and all blocks containing five epochs. Targets appeared in a pseudorandomised sequence, such that within each epoch all eight targets appeared once but across epochs the target sequence was different and participants were unable to predict the next target location. Participants had to perform the movement within a time window of 750 ms, after which the target disappeared. The time cut-off was to ensure the short, ballistic nature of the movement and to prevent online corrections. No endpoint feedback was provided.

At the beginning of each session, participants practised the task by performing 40 baseline trials outside of the scanner (no rotation imposed). Participants then performed the main task inside the scanner, during which they learned to adapt to a stepwise increasing rotation of their movement in case of the adaptation task (Fig. 1A). After the scan, participants performed 144 trials of washout trials (i.e. 3.6 blocks) with no rotation imposed outside of the scanner to probe retention of the previously learned compensatory movement.

### Behavioural data analysis

Cursor movements were analysed on a trial-by-trial basis using in-house software written in Matlab (Mathworks Inc, Natick, USA). The joystick position data (X and Y) was collected at a sample rate of 60 Hz. The kinematic data was filtered with a zero-phase filter with a 25 Hz cut-off and numerically differentiated to determine velocity. Trials that showed premeditated or incomplete movements were excluded from further analysis (mean number of rejected trials: 18.21± 19.3).

For each remaining trial, the angular error (°) was calculated as the oriented angle between a line connecting the starting position with the position of peak velocity of the cursor and the line connecting the starting position with the target where positive values represent a clockwise error (’overshooting’) and negative values indicate a counter-clockwise error (’undershooting’).

#### Behavioural data analysis

Data was grouped into epochs, such that each epoch contained eight consecutive trials and all blocks contained five epochs. To quantify adaptation, we calculated the mean error across all rotation blocks that were included into the analysis, excluding the first epoch of each block respectively {Galea2011}, referred to as adaptation error. To quantify retention, we calculated the mean error in the first washout block, referred to as retention error. The equivalent metrics were also calculated in the control condition, although no rotation was imposed and therefore no adaptation took place.

### MR acquisition

Magnetic Resonance (MR) data were acquired using a 7T MAGNETOM Siemens scanner (Erlangen, Germany) using a 32-channel head coil. Subjects fixated on a crosshair image presented centrally on the screen during resting fMRI acquisition, performed a visuomotor rotation task during MRS acquisition and watched a nature video at all other times.

#### Structural MRI

Structural T1-weighted MRI data were acquired using a magnetisation prepared rapid acquisition gradient echo (MPRAGE) sequence (Repetition Time 2200 ms, TE 2.82 ms, TI 1050 ms, slice thickness 1.0 mm, in-plane resolution 1.0 x 1.0 mm^2^, GRAPPA factor = 4, flip angle = 7 degrees, FOV 192 x 176 mm^2^).

#### MRS

Structural images were used to manually place a voxel over the left precentral knob, which is a landmark for the hand motor representation {Yousry1997}. MRS data were acquired using a semi-LASER sequence {Scheenen2008} (53 transients in each block, TR 6000, TE 36 ms, voxel size 20 x 20 x 20 mm^3^) using VAPOR (variable power RF pulses with optimized relaxation delays) water suppression {Tkac1999}. Shimming of the MRS voxel region used FASTMAP {Gruetter1993}.

Spectra were acquired in 5 1/2 minute MRS-blocks, each corresponding to two Task-blocks., meaning that one MRS-block was acquired at baseline, and six MRS-blocks were acquired during adaptation.

#### Functional MRI acquisition

Resting state fMRI data were acquired before and immediately after the task using a multiband 2 mm isotropic EPI sequence (TR/TE = 3500/28 ms, FOV = 256 x 138 mm^2^, bandwidth =1562 Hz/Px, multi-band acceleration factor 2, voxel dimension = 2 x 2 x 2.5 mm^3^, whole brain, acquisition time = 8:05 min for a total of 132 volumes).

### MR analysis

#### fMRI analysis

Resting-state functional data processing was carried out using FSL (FMRIB’s Software Library, Version 6.00, {Jenkinson2012}).

Standard pre-processing steps were applied: data was motion corrected using MCFLIRT {Jenkinson2002}, slice-timing corrected using Fourier-space time-series phase-shifting, distortion corrected using fieldmaps (B0 unwarping), stripped of non-brain voxels using BET {Smith2002}, grand-mean intensity normalised and highpass temporally filtered (Gaussian-weighted least-squares straight line fitting, with sigma = 50.0s). Probabilistic Independent Component Analysis (ICA), as implemented in MELODIC {Beckmann2004}, was used to manually denoise the data. Data were then registered to the high resolution structural image using boundary-based registration (BBR; implemented in FLIRT {Jenkinson2001} {Jenkinson2002}) and then non-linearly registered to the MNI-152 template and spatially smoothed using a Gaussian kernel of 5mm FWHM.

#### Seed-based connectivity analysis

We quantified functional connectivity between M1 and the cerebellum using a region of interest (ROI)-based approach. Our M1 ROI was defined anatomically and included the anterior bank of the central sulcus as well as the posterior half of the precentral gyrus. It extended from the level of the dorsal surface of the lateral ventricles to the dorsal surface of the brain and from the lateral surface of the brain to the interhemispheric fissure. The cerebellar ROI was defined functionally from a quantitative meta-analysis which identified cerebellar regions involved in adaptation {Hardwick2013}. Specifically, the cerebellar ROI was located in the right cerebellar lobules V-VI. A control ROI for the cerebellum was defined by mirroring the right cerebellar ROI along the midline, such that it was located in the left cerebellar lobules V-VI. M1-Cerebellar Connectivity was quantified by extracting the timeseries of the resting-state activity from the M1 ROI and the cerebellar ROI and correlating the two timeseries. Change was calculated as Post – Pre, with positive values therefore reflecting an increase in connectivity.

#### Network analysis

To replicate the previously reported effect of adaptation increasing cerebellar network-level connectivity (Albert et al, 2009), we ran a group-level ICA with 25 components, as implemented in MELODIC {Beckmann2004}. The cerebellar network of interest was identified by visual inspection and confirmed as the network with the greatest spatial correlation with cerebellar regions active during performance of sensorimotor tasks {Hardwick2013}, as determined by FSL’s fslcc tool for calculating spatial cross-correlation {FSLSmith2004}. To determine the anatomical specificity of any effects we identified a control network which we expected to be unaffected by task condition. We chose the Default Mode Network (DMN) as this is a well defined network that did not spatially overlap with the task activation mask. In order to obtain subject-specific maps, we ran dual regression on both these networks. We then tested for a significant effect of condition on the network using non-parametric permutation inference testing, as implemented in Randomise {Winkler2014}, with a cluster-threshold of 2.3, and p< 0.05, {Woo2012}).

#### MRS analysis

MRS data were pre-processed using in-house scripts and the FID-A toolbox {Simpson2015} according to the steps recommended in {Near2020}. For each session, raw MRS data from all MRS-blocks were concatenated and corrected for frequency and phase shifts. Residual water signal was removed from the water suppressed MRS data using HLSVD {Cabanes2001}. Outlier spectra were identified for each session by calculating a deviation metric for every data point. Data points whose deviation metric exceeded three standard deviations from the mean were removed. To ensure the same number of data points in each MRS-block, these data points were replaced with an interpolated spectrum (maximum 6 (0.016 %) data points per session). Data points were frequency- and phase-corrected and then divided into MRS-blocks. Data was corrected for eddy currents. Neurochemical quantification was performed using LCModel {Provencher2001}. Default LCModel concentration ratio priors were used during fitting. Spectra were then excluded if they had LCModel defined FWHM > 15 Hz or Signal-to-Noise Ratio (SNR) < 40. GABA and Glutamate measurements were excluded if they had CRLB > 50%. Correlations between GABA and other metabolite spectra were < 0.23, indicating that we were able to independently estimate GABA concentration from the MRS spectra (Fig. 3A). To control for differences in voxel tissue composition across subjects, the proportion of Grey Matter (GM), White Matter and CSF in the voxel were quantified from the T1 scan using FMRIB’s Automated Segmentation Tool (FAST) and GM fraction was included as a covariate of no interest when testing for relationships between GABA or Glutamate and another variable {Zhang2001}. All neurochemicals are expressed as a ratio of total creatine (tCr, creatine + phosphocreatine) for internal referencing.

## QUANTIFICATION AND STATISTICAL ANALYSIS

Statistical analyses were conducted using R {RCoreTeam2013} for all analysis apart from the mediation analysis and robust correlation analysis. Mediation analysis was conducted using Statistics Package for the Social Sciences {Hayes2018} {IBMCorp2020}. Robust correlation analysis was performed using the robust correlation toolbox implemented in Matlab {Matlab2019}. To determine whether participants adapted, we used the R package lme4 {Bates2015a} to construct a linear mixed effects model of error during the first and last epoch of all rotation blocks with a fixed main effect of epoch (First, Last). To determine when participants stopped adapting, we calculated post hoc paired-samples t-tests to compare mean error at the start and at the end of each rotation block (Bonferroni-adjusted). Rotation blocks that came after the last block with discernible adaptation were excluded from further analysis. We then tested whether participants retained the compensatory movement. To test for retention, we tested whether the error in the first washout block was significantly different from zero using a one-sample t-test.

Within the period of time where adaptation was detectable, we then tested for significant changes in GABA during adaptation compared to control. To test whether task condition affected GABA over the course of the experiment, we constructed a linear mixed effects model of the connectivity variable of interest. As fixed effects, we entered MRS-block (1, 2, 3, 4, 5, see Fig. 1A), condition (rotation, control) and MRS-block x condition interaction into the model.

We then tested whether task condition affected M1-Cerebellar connectivity by constructing a linear mixed effects model of M1-Cerebellar connectivity. As fixed effects, we entered time (pre, post), condition (rotation, control) and time*condition interaction into the model.

For all linear mixed effects models, we allowed intercepts for different subjects to vary in order to account for covarying residuals within subjects, creating a random intercept model. P-values were obtained by likelihood ratio tests of the full model with the effect in question against the model without the effect in question.

Correlations were assessed using Spearman’s rank correlation coefficients (two-tailed). To determine whether two correlation coefficients were significantly different, Spearman’s coefficients were converted into z-scores using Fisher’s Z-transformation. We then performed one-tailed z-tests on the z-scores given the standard error of each z-score. This method is standard for testing the equality between Pearson’s correlation coefficients, but less commonly used for Spearman’s coefficients. However, simulations show that this practice is justified for Spearman’s coefficients and more robust than using Pearson’s correlation coefficients on non-normal data to test for differences in correlations {Myers2006}.

To ensure the robustness of the observed significant relationship against potential outliers, we re-calculated the correlation coefficients using a MATLAB toolbox for robust correlation methods {Pernet2013}. For this, we first visually inspected scatter plots of the data for a potential non-linear relationship and then tested the normality assumption for the data. Having confirmed the linearity of the relationship and that the data is normally distributed, we tested for potential univariate and bivariate outliers in the data using the three outlier detection methods implemented in the toolbox (box-plot rule, MAD-median rule, S-outliers {Pernet2013}). On the basis of the intersecting results of all three outlier detection methods, which identified no bivariate outliers, but 2 univariate outliers in the GABA data, we chose the percentage-bend correlations as the robust correlation method, as recommended for calculating correlations in the presence of univariate outliers {Wilcox1994} {Pernet2013}. Our data showed inequal variances and therefore the percentage-bend correlation method was interpreted as significant if the percentile bootstrapped 95% confidence interval (CI) of the correlation coefficient excluded zero, as bootstrapped CIs are less sensitive to violations of the homoscedasticity assumption than the traditional t-tests. All results are interpreted taking into account the bootstrapped CIs of the percentage-bend method {Pernet2013}.

### Mediation Analysis

In order to explore the observed correlations between GABA, connectivity change and retention, a mediation analysis was conducted. A mediation analysis tests whether an observed relationship (path c) might be mediated through an indirect pathway (path ab). It is well suited when a variable X is the logical cause of the mediator variable M and if the mediator M in turn is the logical cause of another variable Y. In this study, M1 [GABA] assessed at baseline (X) is the logical cause of connectivity change during adaptation (M) and this connectivity change is the logical cause of later retention of the adapted movement (Y). We therefore performed a simple mediation to analyse whether the relationship between M1 [GABA] (X) and retention (Y) is mediated by Connectivity Change (M). Specifically, we tested whether connectivity change accounts for the link between GABA and retention. The mediation analysis was performed using regression with bootstrapping implemented in the PROCESS macro for SPSS {Hayes2018}. A significant mediation is evident when the bootstrapped 95%-confidence interval of the parameter estimate of the indirect path (ab) does not include zero.

---

**Supplementary Figure 1.**
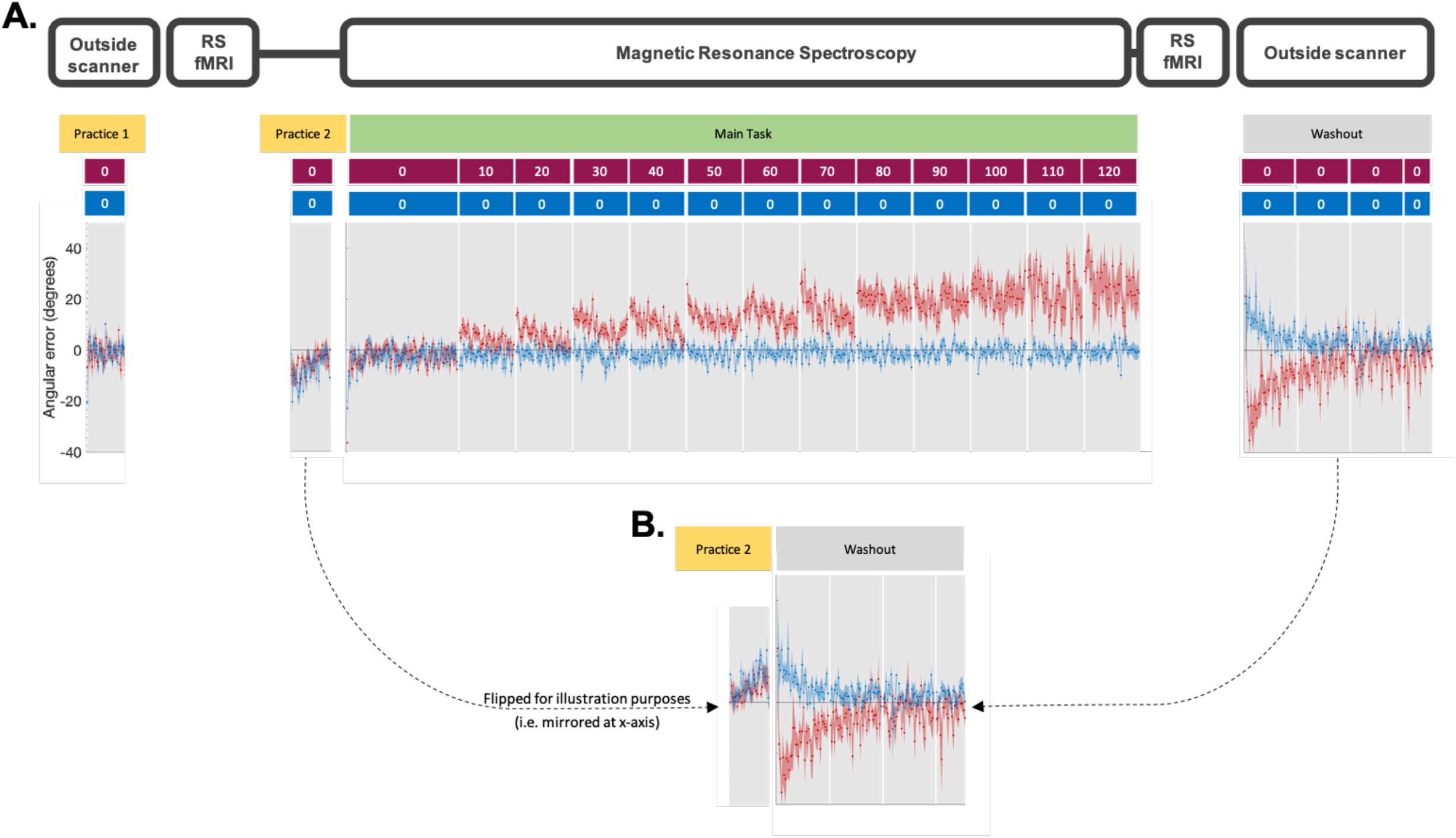
Control condition washout error is driven by movement bias imposed through scanner positioning. **A.** Practice phase inside the scanner reveals unspecific movement bias. Participants practiced the task once outside of the scanner (Practice 1, 32 trials) and once inside the scanner (Practice 2, 32 trials) in every session. As can be seen in practice 2, the positioning inside the scanner imposed an unspecific movement bias that results in a counter clockwise error which reduces to zero within few trials. There is no such movement bias in the practice phase completed outside of the scanner in a seated position (Practice 1), suggesting that this movement bias is driven by placing the participant in a lying-down position in the scanner. **B.** Control condition washout error matches the inverse of the scanner positioning imposed unspecific movement. For illustration purposes, the in-scanner practice has been mirrored at the x-axis. Directly comparing the in-scanner offset with the control condition washout error shows that the two offsets are of similar extent.

## Acknowledgments

The research was supported by the National Institute for Health Research (NIHR), Oxford Biomedical Research Centre and the NIHR Oxford Health Biomedical Research Centre. CJS holds a Sir Henry Dale Fellowship, funded by the Wellcome Trust and the Royal Society (102584/Z/13/Z). HJB holds a Wellcome Principal Research Fellowship (110027/Z/15/Z). JOS is supported by a Sir Henry Dale Fellowship from the Royal Society and the Wellcome Trust (HQR01720). The Wellcome Centre for Integrative Neuroimaging is supported by core funding from the Wellcome Trust (203139/Z/16/Z).

## REFERENCES

Albert, N. B., Robertson, E. M., & Miall, R. C. (2009). The Resting Human Brain and Motor Learning. Curr. Biol. 19(12), 1023–1027. doi:10.1016/j.cub.2009.04.028

Bachtiar, V., Near, J., Johansen-Berg, H., & Stagg, C. J. (2015). Modulation of GABA and resting state functional connectivity by transcranial direct current stimulation. Elife, 4(September 2015), 1–9. doi:10.7554/eLife.08789

Bates, D., Mächler, M., Bolker, B. M., & Walker, S. C. (2015). Fitting linear mixed-effects models using lme4. J. Stat. Softw. 67(1). doi:10.18637/jss.v067.i01. arXiv: 1406.5823

Beckmann, C. F., & Smith, S. M. (2004). Probabilistic Independent Component Analysis for Functional Magnetic Resonance Imaging. IEEE Trans. Med. Imaging, 23 (2), 137–152. doi:10.1109/TMI.2003.822821. arXiv: 978-0-87893-286-3

Cabanes, E., Confort-Gouny, S., Le Fur, Y., Simond, G., & Cozzone, P. J. (2001). Optimization of residual water signal removal by HLSVD on simulated short echo time proton MR spectra of the human brain. J. Magn. Reson. 150 (2), 116–125. doi:10.1006/jmre.2001.2318

Cabral, J., Hugues, E., Sporns, O., & Deco, G. (2011). Role of local network oscillations in resting-state functional connectivity. Neuroimage, 57(1), 130–139. doi:10.1016/j.neuroimage.2011.04.010

Dyke, K., Pépés, S. E., Chen, C., Kim, S., Sigurdsson, H. P., Draper, A., … Jackson, S. R. (2017). Comparing GABA-dependent physiological measures of inhibition with proton magnetic resonance spectroscopy measurement of GABA using ultra-high-field MRI. Neuroimage, 152(October 2016), 360–370. doi:10.1016/j.neuroimage.2017.03.011

Farrant, M., & Nusser, Z. (2005). Variations on an inhibitory theme: Phasic and tonic activation of GABA A receptors. Nat. Rev. Neurosci. 6(3), 215–229. doi:10.1038/nrn1625

Fleury, L., Prablanc, C., & Priot, A. E. (2019). Do prism and other adaptation paradigms really measure the same processes? Cortex, 119, 480–496. doi:10.1016/j.cortex.2019.07.012

Galea, J. M., Vazquez, A., Pasricha, N., Orban De Xivry, J. J., & Celnik, P. (2011). Dissociating the roles of the cerebellum and motor cortex during adaptive learning: The motor cortex retains what the cerebellum learns. Cereb. Cortex, 21(8), 1761–1770. doi:10.1093/cercor/bhq246

Gruetter, R. (1993). Automatic, localized in Vivo adjustment of all first-and second-order shim coils. Magn. Reson. Med. doi:10.1002/mrm.1910290613

Hadipour-Niktarash, A., Lee, C. K., Desmond, J. E., & Shadmehr, R. (2007). Impairment of retention but not acquisition of a visuomotor skill through time-dependent disruption of primary motor cortex. J. Neurosci. 27(49), 13413–13419. doi:10.1523/JNEUROSCI.2570-07.2007

Hardwick, R. M., Rottschy, C., Miall, R. C., & Eickhoff, S. B. (2013). A quantitative metaanalysis and review of motor learning in the human brain. Neuroimage, 67, 283–297. doi:10.1016/j.neuroimage.2012.11.020

Hayes, A. F. (2018a). Introduction to Mediation, Moderation, and Conditional Process Analysis, Second Edition: A Regression-Based Approach.

Hunter, T., Sacco, P., Nitsche, M. A., & Turner, D. L. (2009). Modulation of internal model formation during force field-induced motor learning by anodal transcranial direct current stimulation of primary motor cortex. J. Physiol. 587(12), 2949–2961. doi:10.1113/jphysiol.2009.169284

IBM Corp. (2020). IBM SPSS Statistics for Macintosh, version 27.0.

Jalali, R., Chowdhury, A., Wilson, M., Miall, R. C., & Galea, J. M. (2018). Neural changes associated with cerebellar tDCS studied using MR spectroscopy. Exp. Brain Res. 236(4), 997–1006. doi:10.1007/s00221-018-5170-1

Jayaram, G., Galea, J. M., Bastian, A. J., & Celnik, P. (2011). Human locomotor adaptive learning is proportional to depression of cerebellar excitability. Cereb. Cortex, 21(8), 1901–1909. doi:10.1093/cercor/bhq263

Jenkinson, M., Bannister, P., Brady, M., & Smith, S. (2002). Improved Optimization for the Robust and Accurate Linear Registration and Motion Correction of Brain Images. Neuroimage, 17 (2), 825–841. doi:10.1006/nimg.2002.1132

Jenkinson, M., Beckmann, C. F., Behrens, T. E. J., Woolrich, M. W., & Smith, S. M. (2012). Review FSL. Neuroimage, 62, 782–790. doi:10.1016/j.neuroimage.2011.09.015

Jenkinson, M., & Smith, S. (2001). A global optimisation method for robust affine registration of brain images. Med. Image Anal. 5 (2), 143–156. doi:10.1016/S1361-8415(01)00036-6

Kapogiannis, D., Reiter, D. A., Willette, A. A., & Mattson, M. P. (2013). Posteromedial cortex glutamate and GABA predict intrinsic functional connectivity of the default mode network. Neuroimage, 64(1), 112–119. doi:10.1016/j.neuroimage.2012.09.029

Kolasinski, J., Hinson, E. L., Divanbeighi Zand, A. P., Rizov, A., Emir, U. E., & Stagg, C. J. (2019). The dynamics of cortical GABA in human motor learning. J. Physiol. 597(1), 271–282. doi:10.1113/JP276626

Mawase, F., Bar-Haim, S., & Shmuelof, L. (2017). Formation of long-term locomotor memories is associated with functional connectivity changes in the cerebellar-thalamic-cortical network. J. Neurosci. 37(2), 349–361. doi:10.1523/JNEUROSCI.2733-16.2016

McDougle, S. D., & Taylor, J. A. (2019). Dissociable cognitive strategies for sensorimotor learning. Nat. Commun. 10 (1). doi:10.1038/s41467-018-07941-0

Miall, R. C. [R. C.], & Jenkinson, E. W. (2005). Functional imaging of changes in cerebellar activity related to learning during a novel eye-hand tracking task. Exp. Brain Res. 166(2), 170–183. doi:10.1007/s00221-005-2351-5

Myers, L., & Sirois, M. J. (2014). Spearman Correlation Coefficients, Differences between. Wiley StatsRef Stat. Ref. Online, 1–2. doi:10.1002/9781118445112.stat02802

Near, J., Harris, A. D., Juchem, C., Kreis, R., Marjańska, M., Öz, G., … Gasparovic, C. (2020). Preprocessing, analysis and quantification in single-voxel magnetic resonance spectroscopy: experts’ consensus recommendations. NMR Biomed. (July 2019), 1–23. doi:10.1002/nbm.4257

O’Shea, J., Revol, P., Cousijn, H., Near, J., Petitet, P., Jacquin-Courtois, S., … Rossetti, Y. (2017). Induced sensorimotor cortex plasticity remediates chronic treatment-resistant visual neglect. Elife, 6(September). doi:10.7554/eLife.26602

Oldrati, V., & Schutter, D. J. (2018). Targeting the Human Cerebellum with Transcranial Direct Current Stimulation to Modulate Behavior: a Meta-Analysis. Cerebellum, 17(2), 228–236. doi:10.1007/s12311-017-0877-2

Panouillères, M. T., Joundi, R. A., Brittain, J. S., & Jenkinson, N. (2015). Reversing motor adaptation deficits in the ageing brain using non-invasive stimulation. J. Physiol. 593(16), 3645–3655. doi:10.1113/JP270484

Panouillères, M. T., Miall, R. C. [R. Chris], & Jenkinson, N. (2015). The role of the posterior cerebellum in saccadic adaptation: A transcranial direct current stimulation study. J. Neurosci. 35(14), 5471–5479. doi:10.1523/JNEUROSCI.4064-14.2015

Pernet, C. R., Wilcox, R., & Rousselet, G. A. (2013). Robust correlation analyses: False positive and power validation using a new open source matlab toolbox. Front. Psychol. 3 (JAN), 1–18. doi:10.3389/fpsyg.2012.00606

Petitet, P., Reilly, J. X. O., Gonçalves, A. M., Salvan, P., Kitazawa, S., Johansen-berg, H., & Shea, J. O. (2018). Causal explanation of individual differences in human sensorimotor memory formation, 1–55.

Pinto, A. D., & Chen, R. (2001). Suppression of the motor cortex by magnetic stimulation of the cerebellum. Exp. Brain Res. doi:10.1007/s002210100862

Provencher, S. W. (2001). Automatic quantitation of localized in vivo 1H spectra with LCModel. NMR Biomed. 14(4), 260–264. doi:10.1002/nbm.698

R Development Core Team 3.0.1. (2013). A Language and Environment for Statistical Computing. R Foundation for Statistical Computing. Vienna, Austria. Retrieved from http://www.r-project.org

Richardson, A. G., Overduin, S. A., Valero-Cabré, A., Padoa-Schioppa, C., Pascual-Leone, A., Bizzi, E., & Press, D. Z. (2006). Disruption of primary motor cortex before learning impairs memory of movement dynamics. J. Neurosci. 26 (48), 12466–12470. doi:10.1523/JNEUROSCI.1139-06.2006

Scheenen, T. W., Klomp, D. W., Wijnen, J. P., & Heerschap, A. (2008). Short echo time 1H-MRSI of the human brain at 3T with minimal chemical shift displacement errors using adiabatic refocusing pulses. Magn. Reson. Med. 59(1), 1–6. doi:10.1002/mrm.21302

Semyanov, A., Walker, M. C., Kullmann, D. M., & Silver, R. A. (2004). Tonically active GABAA receptors: Modulating gain and maintaining the tone. Trends Neurosci. 27(5), 262–269. doi:10.1016/j.tins.2004.03.005

Shibata, K., Sasaki, Y., Bang, J. W., Walsh, E. G., Machizawa, M. G., Tamaki, M., … Watanabe, T. (2017). Overlearning hyperstabilizes a skill by rapidly making neurochemical processing inhibitory-dominant. Nat. Neurosci. 20(3), 470–475. doi:10.1038/nn.4490

Shmuel, A., & Leopold, D. A. (2008). Neuronal correlates of spontaneous fluctuations in fMRI signals in monkey visual cortex: Implications for functional connectivity at rest. Hum. Brain Mapp. doi:10.1002/hbm.20580

Simpson, R., Devenyi, G. A., Jezzard, P., Hennessy, T. J., & Near, J. (2017). Advanced processing and simulation of MRS data using the FID appliance (FID-A)—An open source, MATLAB-based toolkit. Magn. Reson. Med. 77(1), 23–33. doi:10.1002/mrm.26091

Smith, J. K., Londono, A., Castillo, M., & Kwock, L. (2002). Proton magnetic resonance spectroscopy of brain-stem lesions. Neuroradiology, 44(10), 825–829. doi:10.1007/s00234-002-0821-z

Smith, M. A., Ghazizadeh, A., & Shadmehr, R. (2006). Interacting adaptive processes with different timescales underlie short-term motor learning. PLoS Biol. 4(6), 1035–1043. doi:10.1063/1.2184639

Smith, S. M., Jenkinson, M., Woolrich, M. W., Beckmann, C. F., Behrens, T. E., Johansen-Berg, H., … Matthews, P. M. (2004). Advances in functional and structural MR image analysis and implementation as FSL. Neuroimage, 23(SUPPL. 1), S208–S219. doi:10.1016/j.neuroimage.2004.07.051

Stagg, C. J., Bachtiar, V., Amadi, U., Gudberg, C. A., Ilie, A. S., Sampaio-Baptista, C., … Johansen-Berg, H. (2014). Local GABA concentration is related to network-level resting functional connectivity. Elife, 2014(3), 1–9. doi:10.7554/eLife.01465

Stagg, C. J., Bachtiar, V., & Johansen-Berg, H. (2011). The role of GABA in human motor learning. Curr. Biol. 21 (6), 480–484. doi:10.1016/j.cub.2011.01.069

Tamaki, M., Wang, Z., Barnes-Diana, T., Guo, D. A., Berard, A. V., Walsh, E., … Sasaki, Y. (2020). Complementary contributions of non-REM and REM sleep to visual learning. Nat. Neurosci. 23(9), 1150–1156. doi:10.1038/s41593-020-0666-y

Taylor, J. A., & Ivry, R. B. (2011). Flexible cognitive strategies during motor learning. PLoS Comput. Biol. 7(3). doi:10.1371/journal.pcbi.1001096. arXiv: e1001096

Taylor, J. A., Krakauer, J. W., & Ivry, R. B. (2014). Explicit and implicit contributions to learning in a sensorimotor adaptation task. J. Neurosci. 34 (8), 3023–3032. doi:10.1523/JNEUROSCI.3619-13.2014. arXiv: NIHMS150003

The Math Works Inc. (2019). Matlab 2019a.

Tkáč, I., Starčuk, Z., Choi, I. Y., & Gruetter, R. (1999). In vivo 1H NMR spectroscopy of rat brain at 1 ms echo time. Magn. Reson. Med. 41(4), 649–656.

Towers, S. K., Gloveli, T., Traub, R. D., Driver, J. E., Engel, D., Fradley, R., … Whittington, M. A. (2004). α5 subunit-containing GABAA receptors affect the dynamic range of mouse hippocampal kainate-induced gamma frequency oscillations in vitro. J. Physiol. doi:10.1113/jphysiol.2004.071191

Ugawa, Y., Uesaka, Y., Terao, Y., Hanajima, R., & Kanazawa, I. (1995). Magnetic stimulation over the cerebellum in humans. Ann. Neurol. 37(6), 703–713. doi:10.1002/ana.410370603

Weightman, M., Brittain, J. S., Punt, D., Miall, R. C., & Jenkinson, N. (2020). Targeted tDCS selectively improves motor adaptation with the proximal and distal upper limb. Brain Stimul. 13(3), 707–716. doi:10.1016/j.brs.2020.02.013

Wilcox, R. R. (1994). The percentage bend correlation coefficient. Psychometrika, 59(4), 601–616. doi:10.1007/BF02294395

Winkler, A. M., Ridgway, G. R., Webster, M. A., Smith, S. M., & Nichols, T. E. (2014). Permutation inference for the general linear model. Neuroimage, 92, 381–397. doi:10.1016/j.neuroimage.2014.01.060

Woo, Krishan, & Wager. (2012). Cluster-extent based thresholding in fMRI analyses. Neuroimage, 100(2), 130–134. doi:10.1016/j.pestbp.2011.02.012.Investigations. arXiv: NIHMS150003

Yousry, T. A., Schmid, U. D., Alkadhi, H., Schmidt, D., Peraud, A., Buettner, A., & Winkler, P. (1997). Localization of the motor hand area to a knob on the precentral gyrus. A new landmark. Brain, 120(1), 141–157. doi:10.1093/brain/120.1.141

Zhang, Y., Brady, M., & Smith, S. (2001). Segmentation of brain MR images through a hidden Markov random field model and the expectation-maximization algorithm. IEEE Trans. Med. Imaging, 20(1), 45–57. doi:10.1109/42.906424

